# Saturated Fat Impairs Circadian Transcriptomics through Histone Modification of Enhancers

**DOI:** 10.1101/2021.02.23.432336

**Authors:** Nicolas J. Pillon, Laura Sardón Puig, Ali Altıntaş, Prasad G. Kamble, Salvador Casaní-Galdón, Brendan M. Gabriel, Romain Barrès, Ana Conesa, Alexander V. Chibalin, Erik Näslund, Anna Krook, Juleen R. Zierath

## Abstract

Obesity and elevated circulating lipids may impair metabolism by disrupting the molecular circadian clock. We tested the hypothesis that lipid-overload may interact with the circadian clock and alter the rhythmicity of gene expression through epigenomic mechanisms in skeletal muscle. Palmitate reprogrammed the circadian transcriptome in myotubes without altering the rhythmic mRNA expression of core clock genes. Genes with enhanced cycling in response to palmitate were associated with post-translational modification of histones. Cycling of histone 3 lysine 27 acetylation (H3K27ac), a marker of active gene enhancers, was modified by palmitate treatment. Chromatin immunoprecipitation and sequencing confirmed that palmitate exposure altered the cycling of DNA regions associated with H3K27ac. Overlap of mRNA and DNA regions associated with H3K27ac and pharmacological inhibition of histone acetyl transferases revealed novel cycling genes associated with lipid exposure of primary human myotubes. Palmitate exposure disrupts transcriptomic rhythmicity and modifies enhancers through changes in histone H3K27 acetylation in a circadian manner. Thus, histone acetylation is responsive to lipid-overload and redirects the circadian chromatin landscape leading to reprogramming of circadian genes in skeletal muscle.

## INTRODUCTION

Obesity is characterized by increased circulating fatty acids and lipid accumulation in central and peripheral tissues, with associated metabolic disturbances and insulin resistance (1). Obesity is also tightly linked to disrupted circadian rhythms – endogenous 24h cycles that allow organisms to anticipate diurnal changes in physiology and behavior (2–5). Circadian misalignment increases circulating levels of glucose, free fatty acids and triglycerides (6), and adversely affects circulating leptin and ghrelin levels (7), which collectively can have deleterious consequences on whole body glucose and energy homeostasis.

Disruption of the circadian rhythms in humans by sleep deprivation or shift-work increases the risk of cardiometabolic diseases, including type 2 diabetes and obesity (8–10). Similarly, simulated chronic jet-lag in mouse models disturbs circadian rhythms and leads to leptin resistance and obesity (11). Moreover, mice expressing a dysfunctional splice variant of the core circadian gene *Clock* are hyperphagic and develop obesity, with systemic alterations in glucose and energy homeostasis (12). Collectively, these results provide evidence to suggest that crosstalk exists between metabolic health, nutritional status, and circadian rhythms. Nevertheless, the relationship between biological clocks and metabolism is complex and bidirectional, and dietary interventions or metabolic diseases can disrupt circadian rhythms.

Circadian rhythms are controlled by transcriptional regulation and post-translational modifications (13, 14). The core clock is composed of cell-autonomous transcription-translation feedback loops, comprised of a CLOCK:BMAL1 heterodimer that transcribes feedback repressors PER, CRY, and NR1D1 (13, 14). The CLOCK:BMAL1 heterodimer can also be coupled to epigenomic mechanisms via histone modifiers, with CLOCK acting as a histone acetyltransferase, thereby altering gene expresison through post-translational modifications (15–20). Intimate links between epigenetic regulation and the circadian clock exist that are likely to contribute to the plasticity of insulin sensitive organs and metabolic control.

Circadian transcription is synergistically regulated by different environmental factors, including energetic states, levels of metabolites, and availability of fuel substrates (21–27). Calorie restriction enhances the amplitude of clock genes and results in an accumulation of histone acetylation (H3K9/K14 and H3K27) at circadian hepatic promoters (28). Long-term high fat diets alter the core clock machinery and clock-controlled genes in mouse tissues (29, 30). Thus, dietary factors reprogram the circadian clock through epigenetic processes, leading to circadian dysregulation of metabolic homeostasis and the onset of metabolic diseases.

Men and women with obesity present altered mRNA expression of core clock genes in skeletal muscle, blood cells and visceral adipose tissue (3–5). Moreover, the altered core clock gene expression in skeletal muscle from men and women with obesity correlates with circulating fatty acid levels (3). In mouse model, diet-induced obesity reprograms circadian gene regulation through rewiring of lipid metabolic pathways (31, 32). Given the link between clock gene expression, lipid metabolism, and metabolic disease, we tested the hypothesis that saturated fatty acids may alter the rhythmicity of gene expression in skeletal muscle. We discovered that the saturated fatty acid palmitate regulates enhancer activity through the regulation of histone H3 lysine K27 acetylation and alters skeletal muscle circadian transcriptomics.

## RESULTS

### Palmitate treatment alters circadian oscillations in primary human myotubes

Primary skeletal muscle cells were synchronized, treated with palmitate or BSA-vehicle, and harvested every 6h (Figure 1A). The RAIN algorithm was used to determine rhythmic oscillations of genes and identified 37% of all transcripts as cycling (Figure 1B). Genes cycling in the control condition, but not after palmitate treatment, were considered repressed by palmitate and represented 20% of all transcripts and over 50% of cycling transcripts (Figure 1B). Conversely, genes not cycling in the control condition, but cycling after palmitate treatment, were induced by palmitate and represented 11% of all transcripts. Genes cycling in both BSA-vehicle and palmitate-treated conditions were considered unaffected by palmitate and only represented 6% of all detected transcripts (Figure 1B). Core clock genes including *BMAL1, CIART, DBP, CRY1, CRY2, NR1D1, NR1D2, PER1, PER2* and *PER3*, were unaffected by palmitate (Figure 1C). Only *CLOCK* and its paralog *NPAS2* were rhythmic in BSA-vehicle-treated, but not in palmitate-treated myotubes (Figure 1C). Thus, palmitate influenced the number of rhythmic genes, with minimal effect on mRNA expression of core clock machinery components.

**Figure 1.**
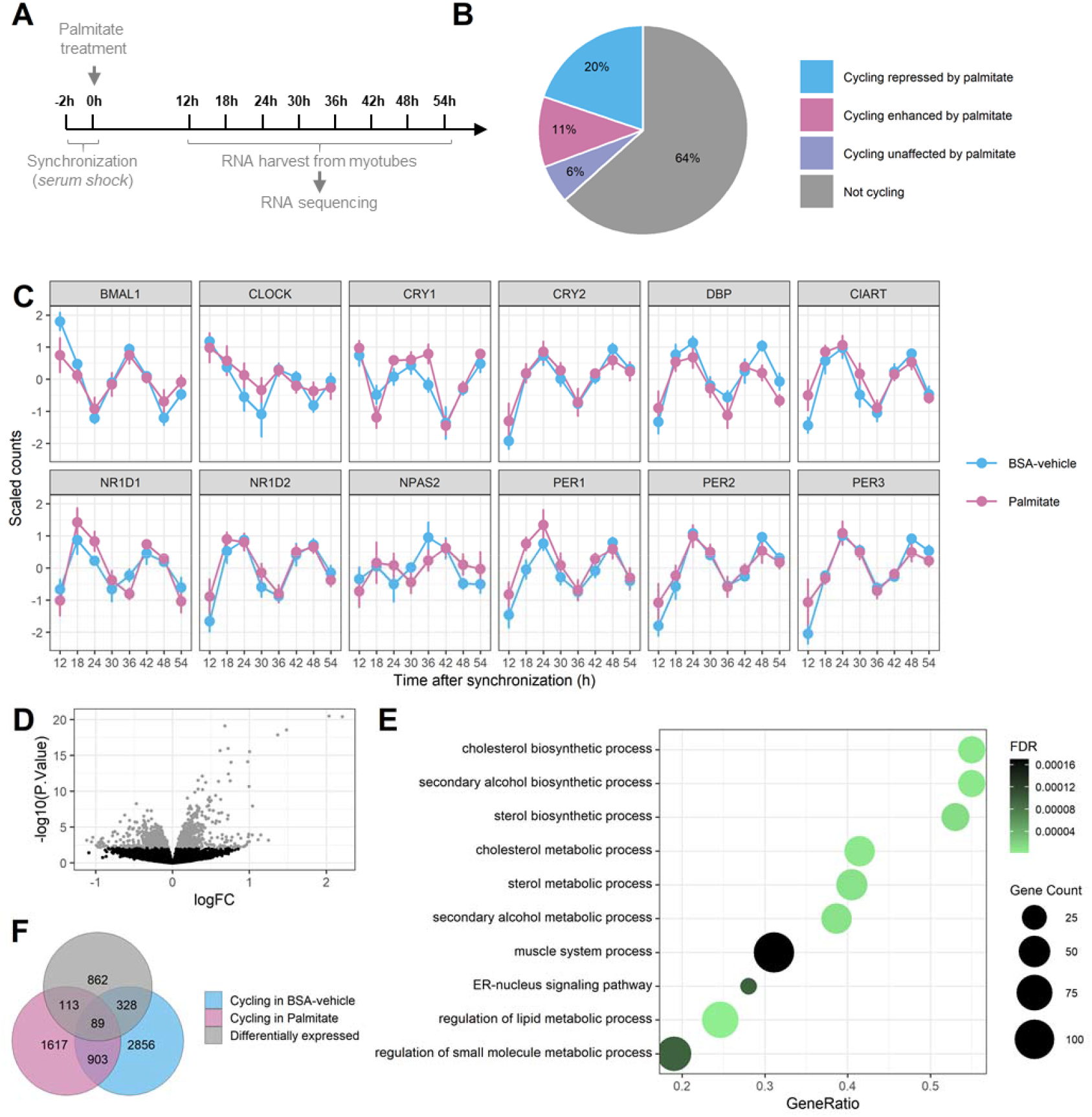
Palmitate alters the pattern of rhythmic transcripts. **A.** Graphic representation of data collection RNA-seq of synchronized primary human skeletal muscle myotubes (n=7) treated with palmitate (0.4 mM) or BSA-vehicle. **B.** Proportion of rhythmic genes (RAIN analysis, FDR<0.1) cycling only in the BSA-condition (repressed by palmitate), only in the palmitate condition (enhanced by palmitate) or in both conditions (unaffected by palmitate). **C.** Core clock genes expression in synchronized myotubes. Data are mean ± SE, n=7. **D.** Differential expression (DE) analysis of rhythmic genes in palmitate-treated myotubes (Limma, FDR<0.1) **E.** Gene set enrichment analysis of gene ontology biological processes (BP) for differentially expressed genes. **F.** Overlap of genes differentially regulated by palmitate (Limma, FDR<0.1) and genes with cycling repressed or enhanced by palmitate (RAIN FDR<0.1).

Changes in gene expression after exposure to saturated fatty acids have been extensively described in multiple model systems (33). To test whether genes where cycling was affected by palmitate overlapped with genes where total mRNA abundance is typically changed in response to fatty acids, differential expression analysis was performed after blocking for the effect of time. Changes in total mRNA abundance after palmitate exposure were observed in 9% of all transcripts (Figure 1D). Palmitate-responsive genes were associated with gene ontology pathways related to lipid metabolism (Figure 1E). Little overlap was observed between genes differentially expressed and genes with dysregulated cycling (Figure 1F), suggesting that an alteration of circadian rhythmicity by palmitate treatment involves mechanisms separate from the canonical lipid metabolic pathways activated by fatty acids.

### Distinct rhythmic gene ontologies in palmitate-treated human myotubes

Gene ontology over-representation analysis for biological processes on the cycling genes unaffected by palmitate showed an enrichment for rhythmic processes and circadian regulation (Figure 2A). Most core clock genes belong to these pathways, confirming the absence of effect of palmitate on the mRNA cycling of the core clock and suggesting that palmitate reprograms circadian transcriptome independently of the core clock machinery. Genes where cycling was repressed by palmitate were associated with pathways involved in transcription and protein targeting to membrane (Figure 2A). Genes where cycling was induced in response to palmitate were annotated to pathways related to post-translational modifications of histones (Figure 2A).

**Figure 2.**
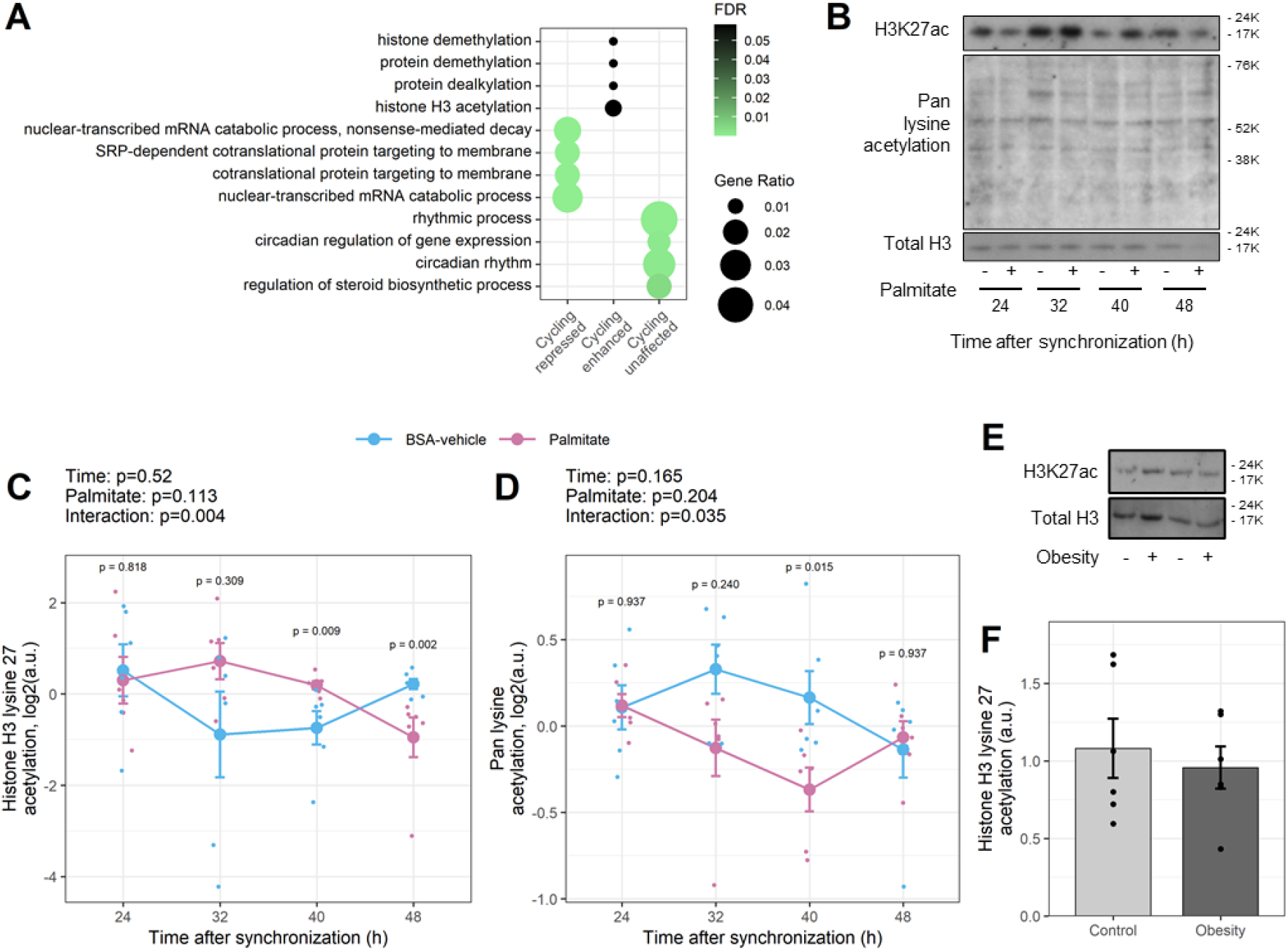
Palmitate alters the circadian acetylation of histone H3 on lysine 27. **A.** Gene ontology overrepresentation analysis of biological processes on genes cycling only in BSA-vehicle, only in palmitate or in both conditions. **B.** Representative immunoblots of histone 3 acetylation from BSA-vehicle- and palmitate-treated myotubes every 8h from 24h to 48h after synchronization. **C-D.** Quantification of lysine 27 acetylation and pan lysine acetylation. Results are mean ± SEM, n=6, paired two-way ANOVA (time, palmitate), followed by pairwise Wilcoxon comparisons. **E-F.** Skeletal muscle biopsies were obtained from men with normal weight (n=6) or obesity (n=6). Protein lysates were collected and relative abundance of total histone H3 and histone H3K27ac was measured. Results are mean ± SEM, n=6.

To explore a potential mechanism for altered temporal regulation of transcription, we investigated whether palmitate treatment altered cyclic histone modifications in synchronized primary human myotubes. Relative abundance of total histone H3, total acetylated lysine, and the marker of active enhancers, histone H3 lysine 27 (H3K27ac), were assessed over a 48h period in BSA-vehicle and palmitate-treated myotubes (Figure 2B). Histone H3 protein abundance was unaffected by time or palmitate treatment, whereas the acetylation of histone 3 on lysine 27 (H3K27ac) was affected by palmitate in a time-dependent manner (RAIN p < 0.05, Figure 2C). Total lysine acetylation exhibited similar palmitate-affected rhythms (RAIN < 0.05), but this regulation was opposite to that of H3K27ac (Figure 2D), suggesting palmitate exposure led to a specific enrichment in histone H3K27ac over total cellular lysine acetylation.

The abundance of H3K27ac was unaltered in *vastus lateralis* skeletal muscle biopsies of from men with obesity versus normal weight (Figure 3E-F), suggesting that palmitate-induced changes in histone acetylation are a consequence of acute fatty acid exposure rather than chronic obesity.

**Figure 3.**
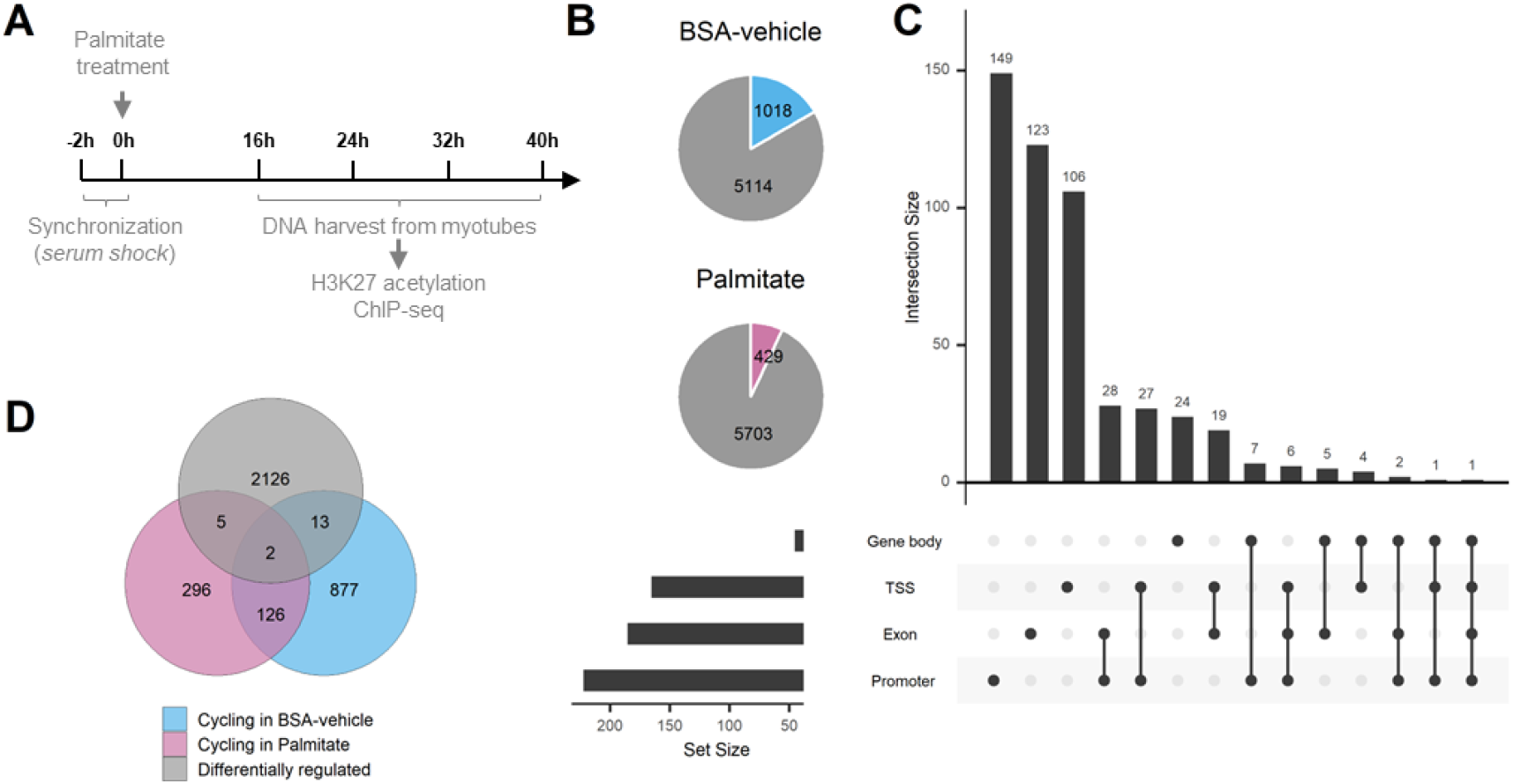
Palmitate treatment reduces the cycling of H3K27 acetylated regions. H3K27ac ChIP-seq of synchronized primary human skeletal muscle myotubes (n=4) treated with palmitate (0.4 mM) or BSA-vehicle. **A.** Graphic representation of data collection **B.** Proportion of rhythmic regions identified (RAIN analysis, FDR<0.1). **C.** Location of acetylated regions in gene body, transcription starting site (TSS), exons or promoter regions of their associated genes. **D.** Differential acetylation analysis compared to regions cycling in BSA-vehicle or palmitate-treated myotubes.

### Palmitate attenuates rhythmic behavior of H3K27 acetylated regions

Genome-wide analysis of histone H3K27ac was performed by chromatin immunoprecipitation (ChIP) and sequencing of synchronized primary human muscle cells treated with palmitate or BSA-vehicle (Figure 3A). We identified more than 127,000 regions with acetylated peaks that were distributed throughout the genome. From these, 6,132 regions were annotated to a gene and used for further analysis. In BSA-vehicle-treated myotubes, we identified 1,018 rhythmic regions, compared to 429 rhythmic regions in palmitate-treated myotubes (Figure 3B). Acetylation regions were predominant in promoter and transcription starting sites, but also present in exons and gene bodies (Figure 3C).

Like the changes observed in mRNA, regions differentially acetylated after palmitate treatment were distinct from the regions with changes in cycling (Figure 4D), suggesting different mechanisms involved in the regulation of cycling compared to simple mRNA transcription. These results suggest that palmitate-induced acetylation of specific H3K27ac enhancer regions is regulated in a circadian manner, and this may be responsible for changes in the rhythmicity of gene expression in response to fatty acids.

**Figure 4.**
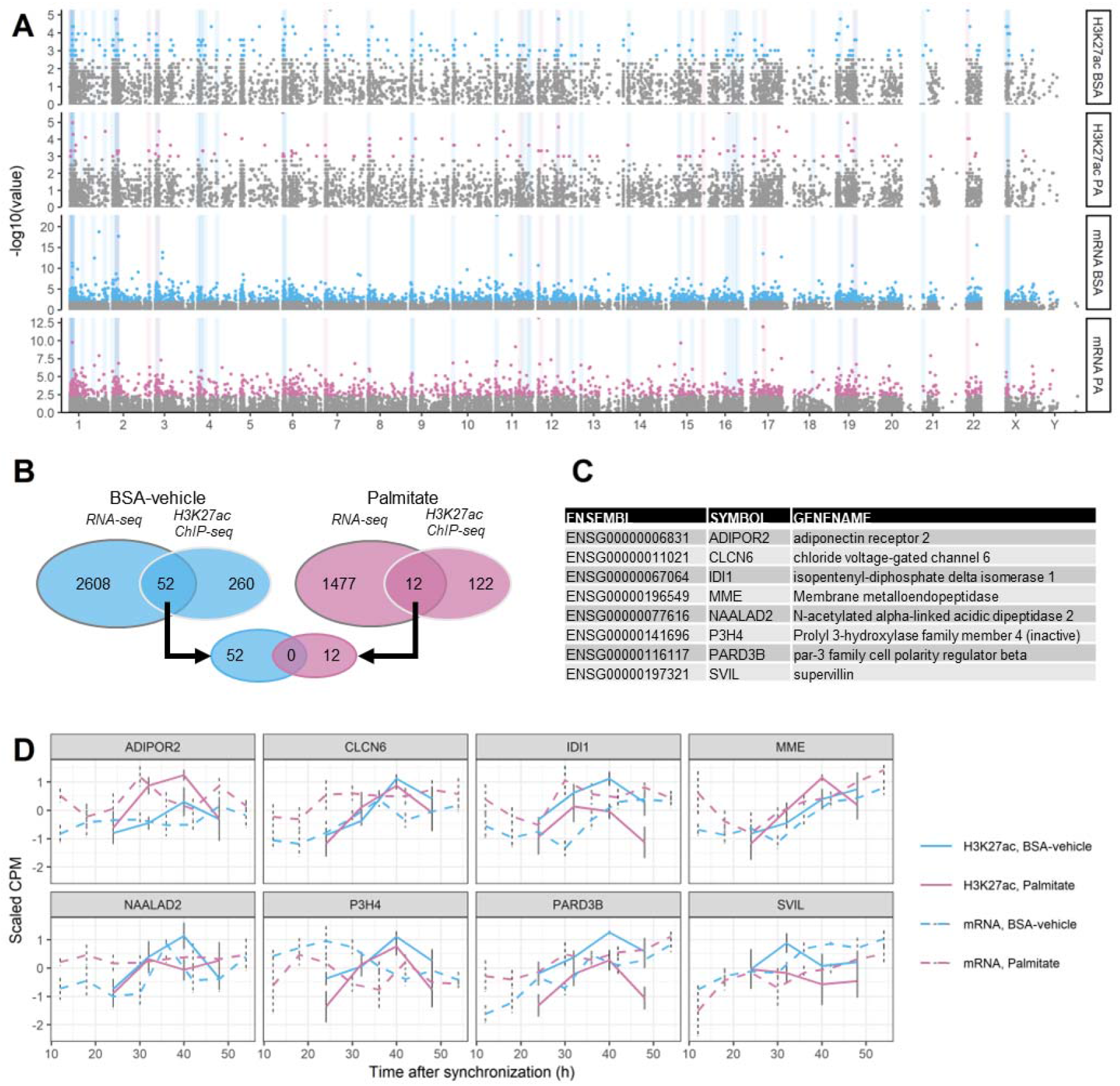
Changes in enhancer rhythmic acetylation and rhythmic transcriptomics in myotubes. **A.** Manhattan plot of cycling genes and regions associated with H3K27ac. From top to bottom: cycling H3K27ac in BSA-vehicle, cycling H3K27ac in palmitate, mRNA cycling in BSA-vehicle, mRNA cycling in palmitate. Grey areas highlight regions of the genome with cycling at both the mRNA and H3K27ac in either BSA-vehicle or palmitate. **B.** Overlap of rhythmic genes (RNA-seq, FDR<0.05) and genes associated to rhythmic regions (H3K27ac ChIP-seq, FDR<0.05) in BSA-vehicle-treated (blue) and palmitate-treated (pink) myotubes. **C-D.** Genes with both cycling mRNA and H3K27ac-associated regions and differentially expressed in myotubes. Data are from the transcriptomics (n=7) and ChIPseq analysis (n=4).

### Changes in rhythmic acetylation of enhancers affect palmitate-responsive genes

Our analysis revealed that palmitate affects the circadian rhythm of many genes, independent of changes in differential expression. Furthermore, we provide evidence that palmitate regulates the acetylation of H3 on lysine 27, a hallmark of enhancers. Thus, we next identified genes with coincident rhythmic mRNA profiles and enhancer regions. Manhattan plots demonstrated that cycling genes and enhancer regions were distributed across the entire genome, with no particular enrichment localized to any specific chromosome (Figure 4A). We identified 52 genes in BSA-vehicle-treated and 12 genes in palmitate-treated myotubes with cycling at both the mRNA and H3K27ac regions (Figure 4B). Of those, only 8 genes were also differentially expressed in myotubes (Figure 4C), confirming that changes in mRNA rhythms can occur independently of changes in total mRNA abundance.

Histone acetyl transferases (HATs) are the main enzymes responsible for acetylation of histones and could therefore be responsible for the changes in rhythmic transcription observed in response to palmitate. Synchronized myotubes were exposed to the histone acetyl transferases inhibitor C646 throughout the palmitate treatment (Figure 5A) to inhibit the acetylation of histone H3K27 (Figure 5B). Histone acetyl transferase inhibition altered the mRNA expression of 5 out of the 8 genes identified previously (Figure 5C). Transcription was either enhanced in response to histone acetyl transferase inhibition (CLCN6, MME, PARD3B) or reduced (P3H4, SVIL), suggesting complex interactions between histone acetylation and the response to fatty acids.

**Figure 5.**
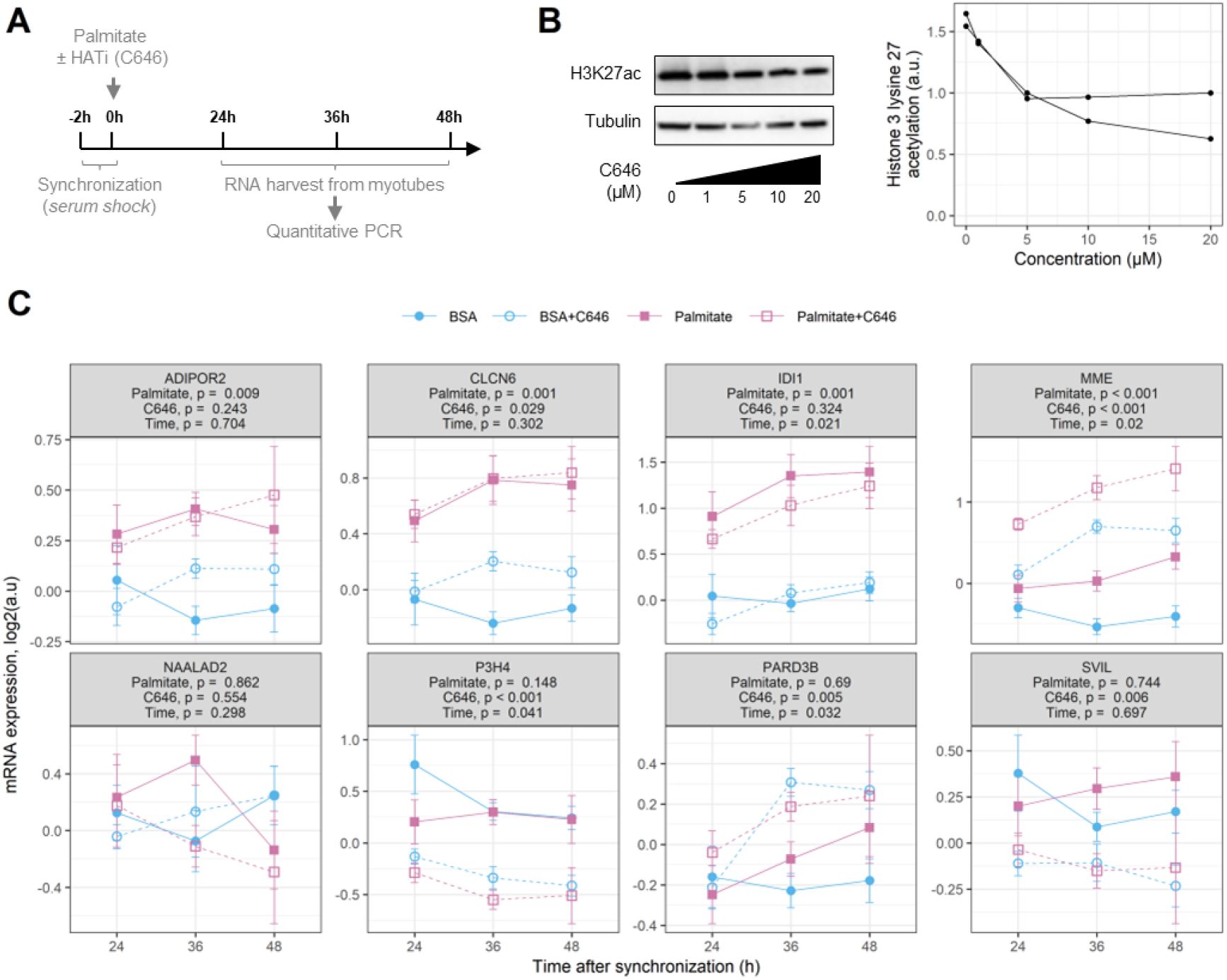
Inhibition of histone acetyl transferases affects mRNA expression of palmitate-responsive genes. **A.** Graphic representation of data collection. **B.** Efficiency of inhibition of histone 3 lysine 27 acetylation by the histone acetyl transferases C646. Representative blots and quantification. **C.** mRNA expression of cycling genes identified as regulated by palmitate in figure 4. Data are mean ± SEM, n=8, results of a 3-way ANOVA (palmitate, C646, time) are presented under the gene names.

## DISCUSSION

Acute saturated fatty acid exposure disrupts rhythmic gene expression in skeletal muscle (3), consistent with fatty acid-induced clock disturbances in liver and adipose tissue (5, 34). Here we show that the saturated fatty acid palmitate triggers transcriptomic and epigenomic changes in skeletal muscle cells that involve alterations of histone acetylation and activation of enhancers leading to disruption of circadian gene expression. Palmitate treatment repressed rhythmicity of genes involved in protein translation and transport, and enhanced rhythmicity in genes involved in histone modifications. Palmitate changed the rhythmicity of enhancers via modification of histone H3K27ac leading to changes in the circadian rhythmicity of selective subsets of genes. Our findings provide insight into mechanisms by which saturated fatty acids alter metabolic homeostasis via circadian misalignment.

Long term palmitate exposure has cytotoxic effects and induces insulin resistance (35, 36). These disturbances coincide with changes in transcription and altered lipid metabolism (34, 37, 38). Increased fatty acid levels promotes lipid oxidation, which is the primary contributor to global acetyl-CoA pool (39). Increased acetyl-CoA levels promote histone acetylation (39), one of the most prominent marks leading to activation of gene expression (13). Here we demonstrate that in skeletal myotubes, palmitate treatment decreased total lysine acetylation, concomitant with changes in rhythmicity and increased histone H3K27ac. Thus, acute exposure to saturated fatty acids imparts post-transcriptional modifications to histone proteins, supporting emerging evidence for lipid-induced epigenomic control of genes important for homeostatic/lipotoxic programs (40).

Histone acetyltransferases (HAT) and histone deacetylases (HDAC) are enzymes that regulate lysine acetylation of proteins, in particular histones. HATs are linked to the circadian control of metabolism (13). We found that HAT inhibition alters palmitate-induced changes in rhythmic gene expression, indicating that saturated fatty acids affect transcriptomic regulation specifically via activation of HAT. Several metabolic sensors act as intermediates to couple circadian rhythms and metabolism. Sirtuins (SIRTs) are NAD^+^-dependent HDACs that sense cellular energy metabolism and whose activity follows a circadian pattern (41). Components of the core clock machinery, CLOCK:BMAL1, coexist with SIRT1 in a chromatin regulatory complex (41). CLOCK can act as a HAT, with specificity for histones H3 and H4 (16), and interact with SIRT1 to regulate acetylation and deacetylation of BMAL1, respectively, which is essential for circadian regulation of gene expression (17). In hepatocytes and adipocytes, palmitate exposure disrupts circadian gene oscillations by preventing BMAL1 deacetylation and activation and interfering with the CLOCK:BMAL1 interaction (34, 42, 43). Thus, palmitate may alter NAD^+^ levels, SIRT1 activity, and CLOCK:BMAL1 action, which may consequently alter histone H3K27ac. Fluctuations of NAD^+^ levels are linked to peripheral clocks (44–46). The extent to which other metabolites and co-factors fluctuate throughout the day to maintain metabolic homeostasis is an emerging topic of research. Nutrient overload associated with high-fat feeding alters tissue-specific metabolomic profiles and leads to circadian misalignment (47). Exercise and nutritional state at different times of the day also influence tissue-specific metabolomic profiles and enrichment of H3K9ac and H3K27ac target genes within myotubes, suggesting that epigenetic modifications of chromatin are influenced by energetic states and fuel substrates (48). The nature of the palmitate-induced intercellular metabolite that governs epigenetic control of the clock machinery requires further interrogation. Nevertheless, our results have physiological implications by linking an oversupply of nutrients in the form of saturated fatty acids to the circadian machinery and the control of metabolism.

We observed circadian oscillations in the levels of histone H3K27ac in primary human muscle cells, supporting the notion that histone modifications are under circadian regulation (20, 28, 29, 49). Palmitate-treatment altered global H3K27ac levels, consistent with evidence that palmitate acetylates enhancer regions regulating lipid metabolism in skeletal muscle and liver cells (50, 51). Changes in histone H3 acetylation are an underlying mechanism regulating rhythmic transcriptional activity in mouse liver (52). However, histone acetylation and specifically histone H3K27ac is likely accompanied by parallel changes in other covalent modifications that also regulate transcription (53). Changes in other histone marks, DNA methylation, mRNA stability and/or post-transcriptional RNA processing may also contribute to the transcript oscillations, even in the absence of enhancer regulation.

Our finding that palmitate treatment increased H3K27ac in cultured myotubes suggests that an acute elevation in circulating lipids may be a contributing factor to this histone modification. Changes in histone H3K27ac in skeletal muscle has been described in the context of aging, with increased expression of genes regulating extracellular matrix structure and organizations (54). We found histone H3K27ac was associated with changes in genes involved in fatty acid metabolism, consistent with earlier reports in pancreas and colon of diet-induced obese mice (51, 55). Conversely, histone H3K27ac was unaltered in skeletal muscle of men with obesity as compared to normal weight. Thus, lipid-induced changes in histone H3K27ac, rather than obesity or insulin resistance may directly contribute to the metabolic dysregulation observed in skeletal muscle.

In summary, the saturated fatty acid palmitate disrupts circadian transcriptomics in primary human myotubes. Our results provide a link between nutrient overload, disruptions of circadian rhythms, and metabolic pathways. Increased histone H3K27ac in palmitate-treated primary human myotubes suggests a specific role for this epigenetic mark in the transcriptional changes that occur in peripheral tissues in response to lipid-overload. Disruption of circadian rhythms in skeletal muscle due to lipid overload may lead to epigenomic changes that influence metabolism. Thus, dietary, or therapeutic modulation of lipid levels, a cornerstone in the treatment of metabolic disorders, may prevent circadian misalignment in peripheral tissues.

## METHODS

### Subjects

*Vastus lateralis* muscle biopsies were obtained from 7 healthy men to establish primary skeletal muscle cell cultures to determine the effects of palmitate on circadian transcriptomics, histone H3 protein abundance, and histone H3 lysine 27 acetylation (H3K27ac). *Vastus lateralis* muscle biopsies were obtained from a cohort of men with normal weight (n=6) or obesity (n=6) as reported earlier (56) and a portion of the biopsy was processed for immunoblot analysis of H3 protein abundance and H3K27ac. Clinical characteristics of the men with normal weight or obesity are presented in Table 1. Studies were approved by the regional ethics committee of Stockholm and conducted in accordance with the Declaration of Helsinki (2012/1955-31/1, 2013/647-31/3, 2012/1047-31/2 and 2016/355-31/4). Participants gave informed consent prior to enrolment.

**Table 1.** Clinical Characteristics of the Study Participants.

|  | **Normal weight** | **Obese** |
| --- | --- | --- |
|  | **Men (n=6)** | **Men (n=6)** |
| Age (yr) | 53 ± 3 | 48 ± 2 |
| Body weight (kg) | 79.8 ± 2.6 | 103.3 ± 1.6* |
| BMI (kg/m<sup>2</sup>) | 24.2 ± 0.3 | 31.1 ± 0.7* |
| fP-Glucose (mmol/L) | 5.4 ± 0.2 | 6.5 ± 0.3 |
| fP-Insulin (pmol/L) | 39.1 ± 7.2 | 116.3 ± 26.8* |
| HOMA-IR | 1.3 ± 0.2 | 4.3 ± 1.2* |
| fP-Cholesterol (mmol/L) |  |  |
| Total | 4.9 ± 0.2 | 5.2 ± 0.5 |
| LDL | 3.3 ± 0.2 | 2.9 ± 0.5 |
| HDL | 1.3 ± 0.05 | 1.6 ± 0.3 |
| fP-Triglycerides (mmol/L) | 0.9 ± 0.1 | 1.8 ± 0.3* |

Results are mean ± SEM for normal weight men and men with obesity. Differences between normal weight and men with obesity were determined using unpaired Student's *t*-test. \*P<0.05 versus normal weight.

### Primary human skeletal muscle cell cultures

Primary myoblasts were grown in DMEM/F12+Glutamax with 16% FBS and 1% Antibiotic-Antimycotic. Cells were regularly tested for mycoplasma contamination by PCR. At 80% confluence, myoblasts were differentiated into myotubes by culturing in fusion medium consisting of 76% DMEM Glutamax with 25 mM glucose, 20% M199 (5.5 mM), 2% HEPES, and 1% Antibiotic-Antimycotic (100X), with 0.03 μg/mL zinc sulfate and 1.4 mg/mL Vitamin B12. Apo-transferrin (100 μg/mL) and insulin (0.286 IU/mL) was added to the fusion medium. After 4-5 days, the medium was switched to the same medium without apotransferrin or insulin, with 2% FBS (post-fusion media) and the cultures were continued for 3-5 days.

### Palmitate treatment

Palmitate stock solution (200 mM) (C16, Sigma P9767) was prepared in 50% ethanol and then diluted 25 times in a 10.5% BSA solution. BSA (Sigma A8806) in serum-free essential alpha medium (αMEM) was used as a carrier and control. Myotubes were incubated in 5.5 mM glucose media for 22h prior to the experiments. Myotubes were synchronized by serum shock (50% FBS, 2h), washed with PBS, and incubated in 5.5 mM glucose medium containing palmitate (0.4 mM) or BSA (vehicle). Cultures were collected every 6h for mRNA analysis and every 8h for DNA and immunoblot analysis, starting from 12h to 54h after synchronization.

### Immunoblot analysis

Myotube cultures were lysed in ice cold buffer A (1% Protease Inhibitor Cocktail, 137 mmol/L NaCl, 2.7 mM KCl, 1 mM MgCl_2_, 5 mM Na_4_P_2_O_7_, 0.5 mM Na_3_VO_4_, 1% Triton X-100, 10% glycerol, 20 mM Tris, 10 mM NaF, 1 mM EDTA, and 0.2 mM phenylmethylsulfonyl fluoride, pH 7.8), followed by end-over-end rotation for 60 min (4°C) and centrifugation at 12,000g for 15 min (4°C). Skeletal muscle biopsies were pulverized in liquid nitrogen and lysed in homogenization buffer A supplemented with 0.5% of NP-40 and 0.02% of SDS, followed by end-over-end rotation for 60 min (4°C) and centrifugation at 3,000g for 10 min (4°C). Protein concentration was determined using a Pierce BCA protein assay kit (#23225, Thermo Fischer Scientific). Samples were prepared for SDS-PAGE with Laemmli buffer, separated on Criterion XT Bis-Tris 4-12% gradient gels (Bio-Rad, Hercules, CA) and transferred to PVDF membranes (Merck). Ponceau staining was performed, and the results were normalized to the total amount of protein per lane. Western blot was performed using primary antibodies (1:1000 concentration) in tris□buffered saline (TBS) containing 0.1% bovine serum albumin (BSA) and 0.1% NaN_3_. Bionordika antibodies for Acetyl-Histone H3 Lysine 27 (#8173S), Acetylated-Lysine (#9441S) and Histone H3 (#4499T) were used. Cell Signaling Technology antibodies for phospho-ACC (Ser79) (#3661), total ACC (#3676), phospho-AMPKα (Thr172) (#2534), and total AMPK (#32532) were used. A mouse monoclonal GAPDH (#sc-47724, Santa Cruz Biotechnology) was used as a loading control. Species□appropriate horseradish peroxidase conjugated secondary antibodies were used at a concentration of 1:25,000 in 5% skimmed milk in TBS Tween. Proteins were visualized by chemiluminescence (#RPN2232 ECL and #RPN2235 ECL select Western blotting detection reagent – GE Healthcare, Little Chalfont, U.K.) and quantified using ImageLab software v. 5.2.1 (BioRad).

### RNA extraction and RNA sequencing

RNA sequencing from myotube cultures was performed as described previously (57). Briefly, RNA was extracted with Trizol Reagent and miRNAeasy kit (Qiagen; Cat #217004) and processed using the Illumina TruSeq Stranded Total RNA with Ribo-Zero Gold protocol (Illumina). Ribosomal RNA was removed and RNA sample fragmented and subjected to first-strand cDNA synthesis. cDNA was subjected to AMPure beads (Beckman Coulter) and adenylated to prime for adapter ligation followed by PCR amplification. Single-end sequencing was performed on the X Ten platform (Illumina) at the Beijing Genomics Institute (BGI; Hong Kong, China). RNA-seq reads 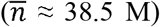 from FASTQ files were qualitytrimmed using Trim_Galore (v0.4.3) and aligned using STAR (v2.5.3a) (58) with Ensembl human annotation (GRCh38, release 92). Gene features were counted using featureCounts (59) from subread (v1.5.2) package and analyzed with edgeR (60). The logCPM (count per million) values for each gene were calculated using limma’s *voom* function while correcting for batch effect from participants using *duplicateCorrelation* function (61).

### H3K27 acetylation chromatin immunoprecipitation and sequencing (ChIP-seq)

Acetylated H3K27 ChIP-sequencing was performed as described (50). Myotubes were cross-linked in 1% formaldehyde in PBS for 10 min at room temperature followed by quenching with glycine (0.125 M). Cells were washed with PBS and harvested in 1 ml SDS Buffer (50 mM Tris-HCl (pH 8), 100 mM NaCl, 5 mM EDTA (pH 8.0), 0.2% NaN3, 0.5% SDS, 0.5 mM phenylmethylsulfonyl fluoride) and subjected to centrifugation (6 min at 250 x g). Pelleted nuclei were lysed in 1.5 ml ice-cold IP Buffer (67 mM Tris-HCl (pH 8), 100 mM NaCl, 5 mM EDTA (pH 8.0), 0.2% NaN3, 0.33% SDS, 1,67% Triton X-100, 0.5 mM phenylmethylsulfonyl fluoride) and sonicated (Diagenode, Biorupter) to an average length of 200–500 bp. Before starting the ChIP experiment, chromatin was cleared by centrifugation for 30 min at 20,000 x g. For each ChIP, 2-10 μg DNA was combined with 2.5 μg antibody for H3K27ac (Ab4729), and incubated with rotation at 4°C for 16 h. Immunoprecipitation was performed by incubation with Protein G Sepharose beads (GE healthcare) for 4 h followed by three washes with low-salt buffer (20 mM TrisHCl (pH 8.0), 2 mM EDTA (pH 8.0), 1% Triton X-100, 0.1% SDS, 150 mM NaCl) and two washes with high-salt buffer (20 mM Tris-HCl (pH 8.0), 2 mM EDTA (pH 8.0), 1% Triton X-100, 0.1% SDS, 500 mM NaCl). Chromatin was de-cross-linked in 120 μl 1% SDS and 0.1 M NaHCO_3_ for 6 h at 65°C, and DNA was subsequently purified using Qiagen MinElute PCR purification kit. For library preparation and sequencing, 3-10 ng of immunoprecipitated DNA was used to generate adapter-ligated DNA libraries using the NEBNext^®^ Ultra DNA library kit for Illumina (New England Biolabs, E7370L) and indexed multiplex primers for Illumina sequencing (New England Biolabs, E7335). The PCR cycle number for each library amplification was optimized by running 10% of the library DNA in a real-time PCR reaction using Brilliant III Ultra-fast SYBR Green QPCR Master Mix (AH Diagnostic) and a C1000 Thermal cycler (Bio-Rad). DNA libraries were sequenced on a HiSeq2000 by 50□bp single□end sequencing at the National High Throughput Sequencing Centre (University of Copenhagen, Denmark).

### Processing of H3K27ac ChIP-seq data

H3K27ac ChIP-Seq samples quality was assessed using FASTQC software. One sample was of poor quality and had to be excluded. Quality was trimmed with trimmomatic (v0.32) (62) on the same parameters to clip adapters to remove low quality sequences using a minimum quality of 30, a sliding window of 5 and a minimum length of 40 bp. Reads were mapped to *H. sapiens* reference genome using Bowtie2 (v2.3.2-foss-2016b) aligner (63). Duplicate removal was performed using samtools (v1.5-foss-2016b) (64) and then BAM files were subjected to peak count using MACS2 software (v2.1.0) (65) for broad peaks using a q-value of 0.01 and a shift of 147 bp. Called peaks were quantified with featureCounts (v1.6.3) (59) and associated to closest genes using RGmatch software (66), mapping regions to the transcriptomic start site and the promoter of the first exon. The resulting peak matrix was Reads Per Kilobase of transcript, per Million mapped reads (RPKM) normalized and batch corrected using ComBat (67) to correct for sequencing lane bias.

### Bioinformatic analysis

Time series samples were analyzed with the R Package RAIN (68) to capture the rhythmic oscillations and determine peak times of the gene expression and acetylated regions. Rhythmicity was determined based on a 24h longitudinal period. Genes and H3K27 acetylated regions were considered rhythmic when FDR<0.1. Limma R package (61) was used to determine differential gene expression and histone acetylation. The effect of time was blocked using a linear function to study the independent effect of palmitate. The effect of palmitate and obesity was considered significant when FDR<0.1. Enrichment of functional clusters was performed on significant genes (FDR<0.1) using clusterProfiler (69).

### Quantitative PCR

Cultured muscle cells were lysed and RNA was extracted using the E.Z.N.A Total RNA Kit (Omega Bio-tek). All equipment, software and reagents for performing the reverse transcription and qPCR were from. cDNA synthesis was performed from ~0.5 μg of RNA using the High Capacity cDNA Reverse Transcription kit (Thermo Fisher Scientific). qPCR was performed on a Viia7 system with TaqMan Fast Universal PCR Master Mix and predesigned TaqMan probes (Thermo Fisher Scientific).

### Statistics

Statistical analyses were performed using R version 4.2.0. Normality was tested using the Shapiro-Wilk’s normality test and equality of variances tested with Levene’s test. Tukey transformation was used when required to run 2-way and 3-way ANOVA to determine the overall effect of palmitate, time and inhibitor in western blot and qPCR data. Student’s unpaired *t*-test was performed to determine the effect of obesity on total histone H3 and histone H3K27ac in skeletal muscle biopsies. P < 0.05 was considered significant.

## Abbreviations

BMAL1: also known as aryl hydrocarbon receptor nuclear translocator-like protein [ARNTL]
CIART: Circadian Associated Repressor of Transcription
CLOCK: Circadian Locomotor Output Cycles Kaput
CRY: Cryptochrome
DBP: D site of albumin promoter (albumin D-box) binding protein
H3K27ac: Histone H3 Lysine 27 acetylation
HPRT: hypoxanthine guanine phosphoribosyl transferase
NR1D1: Nuclear Receptor Subfamily 1 Group D Member 1, also known as REVERBα
NR1D2: Nuclear Receptor Subfamily 1 Group D Member 2, also known as REVERBβ
PER: Period
TBP: TATA box binding protein
TEAD1: Transcriptional enhancer factor TEF-1
18S: 18S ribosomal RNA

## DATA AVAILABILITY

The raw and processed files for the RNAseq and the ChIPseq experiments have been deposited on the GEO repository under accession numbers GSE205424 and GSE205677.

## ACKNOWLEDGEMENTS

The authors are supported by grants from the Novo Nordisk Foundation (NNF14OC0011493, NNF17OC0030088, NNF21SA0072747), Swedish Diabetes Foundation (DIA2018-357, DIA2018-336), Swedish Research Council (2015-00165, 2018-02389), KID-funding (2-3591/2014), the Strategic Research Program in Diabetes at Karolinska Institutet (2009-1068), Marie Skłodowska-Curie Actions (European Commission, 704978), EFSD European Research programme on New Targets for Type 2 Diabetes and the Stockholm County Council (SLL20150517, SLL20170159, SLL20190173). Additional support was received from the Novo Nordisk Foundation Center for Basic Metabolic Research at the University of Copenhagen (NNF18CC0034900).

## AUTHOR CONTRIBUTIONS

NJP, LSP, AA, BMG, RB, AK and JRZ devised the study concept and design.

LSP, BMG and NJP performed the muscle culture experiments.

LSP and NJP performed the sample preparation for RNAseq and ChIP Seq analysis.

LSP and AVC performed the Western Blot analysis.

RB and AA processed the RNA-seq measurements.

LSP, SCG, AC, performed the bioinformatic analysis of the ChIP-seq.

EN measured the anthropometrics and collected the skeletal muscle biopsy material.

NJP, LSP, AK and JRZ. drafted the manuscript. All authors critically revised the manuscript for important intellectual content.

